# A high throughput single molecule platform to study DNA supercoiling effect on protein-DNA interactions

**DOI:** 10.1101/2024.10.24.620099

**Authors:** Huijin Lee, Jihee Hwang, Fahad Rashid, James A. London, Richard Fishel, James M. Berger, Sua Myong, Taekjip Ha

## Abstract

DNA supercoiling significantly influences DNA metabolic pathways. To examine its impact on DNA-protein interactions at the single-molecule level, we developed a highly efficient and reliable protocol to modify plasmid DNA at specific sites, allowing us to label plasmids with fluorophores and biotin. We then induced negative and positive supercoiling in these plasmids using gyrase and reverse gyrase, respectively. Comparing supercoiled DNA with relaxed circular DNA, we assessed the effects of supercoiling on CRISPR-Cas9 and mismatch repair protein MutS. We found that negative DNA supercoiling exacerbates off-target effects in DNA unwinding by Cas9. For MutS, we observed both negative and positive DNA supercoiling enhances the binding interaction between MutS and a mismatched base pair but does not affect the rate of ATP-induced sliding clamp formation. These findings not only underscore the versatility of our protocol but also opens new avenues for exploring the intricate dynamics of protein-DNA interactions under the influences of supercoiling.

## INTRODUCTION

DNA supercoiling, the twisting and winding of the double helix structure beyond its relaxed state, stands as a fundamental mechanism for regulating numerous biological processes (1, 2). From the compact packaging of genetic material within the confines of the cell nucleus to the dynamic modulation of gene activity (3), DNA supercoiling emerges as a versatile molecular switch, controlling vital cellular functions (4). DNA supercoiling regulates DNA accessibility (5), replication and transcription, profoundly impacting gene expression and genome stability. For example, RNA polymerase (RNAP) generates positive supercoiling ahead of the transcription machinery and negative supercoiling behind it (6–8). These accumulated supercoiling states need to be resolved by topoisomerases, and failure to do so can lead to transcriptional arrest (9). Similarly, the replisome generates positive supercoiling in front of it (10–12) and topoisomerases are needed for proper elongation by the replisome and chromosome segregation during cell division (13, 14).

To study how DNA supercoiling affects protein behavior, various techniques have been employed. Traditional methods like electrophoresis differentiate DNA molecules based on their migration speed, which vary according to their supercoiling status (15, 16). However, electrophoresis does not allow real-time observation of functionally relevant dynamics. Single-molecule techniques provide additional insights by allowing precise control of DNA supercoiling while enabling the observation of protein-DNA dynamics in real time. In magnetic tweezers, a single linear double stranded DNA molecule is tethered at both ends, with one extremity linked to a paramagnetic bead surface and the other anchored to the glass surface of a flow cell (17, 18). Tethering to each surface is achieved through multiple bonds, preventing the DNA from swiveling around the attachment points. By rotating the bead through manipulating a magnet above the sample chamber, magnetic tweezers can control torque and twist on DNA in real time (19). Magnetic tweezers allow for precise application of force and torque, making it possible to study the dynamic behavior of DNA-protein interactions under varying supercoiling conditions (5, 20, 21). Additionally, by attaching a fluorescent rotor bead on the side of DNA, it is possible to measure the torque and twist on DNA (22). As the DNA swivels, the rotor bead also swivels, and its associated rotation is observed through fluorescence imaging. This method allows for real-time observation of the rotational dynamics of DNA (23). Optical tweezers offer higher temporal resolution compared to magnetic tweezers, making them suitable for studying fast dynamics (24). Typically, optical tweezers generally do not have the ability to control DNA supercoiling with two notable exceptions accomplished using a birefringent particle instead of a bead (25) or using DNA overstretching followed by strand/bead untethering and re-tethering (26). However, optical tweezers can only manipulate one molecule at a time when introducing supercoiling in DNA, resulting in inherent low throughput.

To address these limitations, we introduce a single-molecule approach to study the supercoiling effect on protein-DNA interactions with high throughput. By implementing single molecule total internal reflection fluorescence (smTIRF) microscopy and utilizing an enzyme-based internal DNA modification method (27), we established a Fluorescence Resonance Energy Transfer (FRET)-based high-throughput assay. This platform allowed us to investigate the effects of DNA supercoiling on two different systems: CRISPR-Cas9 and DNA mismatch repair (MMR). In CRISPR-Cas9 system, we monitored real-time DNA unwinding kinetics as a function of supercoiling in the presence of mismatch sequences. Our results suggest that DNA negative supercoiling enhances DNA unwinding by the nuclease-deficient Cas9 (dCas9) in complex with guide RNA (gRNA) (dCas9-gRNA), in the presence of mismatches either in the protospacer adjacent motif (PAM)-proximal or distal regions. Furthermore, we observed that the dynamics of MutS, a protein that recognizes a mismatch to initiate MMR, depend on DNA supercoiling states. We found that the initial binding between MutS and a mismatched base pair can be enhanced by DNA supercoiling in either direction.

## MATERIAL AND METHODS

### Cloning

Plasmid pUC19HL_MutS and pUC19HL_CRI were prepared by introducing two BbvCI nick sites and one BsmI restriction site to plasmid pUC19. A BsmI site was introduced into the plasmid by site-directed mutagenesis using the Q5 Site-Directed Mutagenesis Kit (NEB, E0554), following the manufacturer’s protocol. The primers used were 5’-ATGCCAAAATCCCTTAACGTG-3’ and 5’-TCATGAGATTATCAAAAAGGATC-3’. Vector DNA was prepared by digestion with EcoRI (NEB, R3101S, 2 unit/μg DNA) and HindIII (NEB, R3104S, 2 unit/μg DNA) of Plasmid pUC19 (NEB, N3041S) supplemented with rCutSmart Buffer (NEB, B6004S) for 15 min at 37 °C. The DNA was purified using PCR spin column (Qiagen, 28104). Insert DNA for pUC19HL_MutS was prepared by annealing two 5’-phosphorylated oligonucleotides 5’AGCTTCCTCAGCTTAATACGACTCACTATAGGCCAATACAAGAGCTTCATCCTCAGCG3’ and 5’AATTCGCTGAGGATGAAGCTCTTGTATTGGCCTATAGTGAGTCGTATTAAGCTGAGGA3’. Insert DNA for pUC19HL_CRI was prepared by annealing two 5’-phosphorylated oligonucleotides 5’ AGCTTCCTCAGCCCAGCGTCTCATCTTTATACATCAGCAGAGATTTCTGCTGTGCAACCTCAGCG3’ and 5’AATTCGCTGAGGTTGCACAGCAGAAATCTCTGCTGATGTATAAAGATGAGACGCTGGGCTGAGG A3’. The vector DNA and insert DNA were then incubated with T4 DNA ligase at room temperature for 10 minutes, followed by inactivation at 60 °C for 10 minutes. The mixture was added to competent DH5α *E. coli* cells (NEB, C2988J) and incubated at 42 °C for 30 seconds for transformation. Transformed cells were cultured in 1 mL of LB broth and plated on LB agar plates supplemented with ampicillin. Several colonies were picked and grown in LB broth. Plasmids from each colony were extracted using a mini prep kit (Omega, D6942). The sequence was confirmed by Sanger sequencing (GENEWIZ from Azenta Life Sciences).

### Labeled oligonucleotides preparation

All DNA oligonucleotides were purchased from Integrated DNA Technologies (IDT). A thymine modified with an amine group through a C6 linker (amino-dT) was used for labeling. All oligonucleotides sequence and the location of amino modification and biotin are shown in Supplementary Table S1. The nucleotide (5 nmol) and sulfo-Cy3 NHS ester (Lumiprobe, 13320) or sulfo-Cy5 NHS ester (Lumiprobe, 11320) (250 nmol) were mixed in 0.1 M NaHCO_3_ solution. The mixture was incubated at 40 °C overnight. The labeled nucleotides were precipitated by incubating with 3 times volume of ethanol at -80 °C for 1 hour. The mixture was centrifuged at 20000 g at 4 °C for 40 min. The remaining dyes were removed by carefully removing the supernatant and washing the pellet with 70% ethanol solution. The residual ethanol was dried in the air and the DNA pellet was suspended in nuclease-free water.

### Strand replacement

Plasmid DNA was purified using QIAGEN Plasmid Plus Maxi kit (QIAGEN, 12963). Plasmid DNA (5 µg) and labelled single-stranded oligo (50-fold excess relative to the DNA molarity) were mixed with either Nt.BbvCI (NEB, R0632) or Nb.BbvCI (NEB, R0631) (1.75 µL, 17.5 units) and rCutSmart buffer (supplied with enzyme) (5 µL) in a total volume of 50 µL. The mixture was incubated at 37 °C for 3 hours and then 95 °C for 5 min. The temperature decreased to 25 °C slowly (1 °C/min). The extra single stranded oligo was removed using 50K amicon (Millipore sigma, UFC5050BK). The purified plasmid DNA (5 µg) was incubated with T4 DNA ligase (NEB, M0202) (2.5 µL, 1000 units) and T4 ligase reaction buffer (supplied with enzyme) (7.5 µL) in a total volume of 75 µL overnight at RT. After the incubation, DNA was purified using PCR spin columns (Qiagen, 28104). The second strand replacement was performed in the same manner as the first strand replacement, but the Amicon purification step was omitted. After annealing, the mixture was incubated with T4 DNA ligase (NEB, M0202) (2.5 µL, 1000 units), supplemented with DTT (1 mM) and ATP (1 mM) in a total volume of 75 µL overnight at RT. The mixture was treated with T5 exonuclease (NEB, M0663) (5 µL, 50 units) for 30 min at 37 °C. After the reaction, DNA was purified using PCR spin columns (Qiagen, 28104). DNA concentration and purity (A260/A230 and A260/A280) were confirmed by Nanodrop. The plasmids at each step were confirmed by 1% agarose gel.

### Gyrase preparation

Full-length GyrA and GyrB were cloned into pRSF vector with double His tag and SUMO tag at the N-terminus. The GyrA and GyrB constructs were transformed into Rosetta™(DE3)pLysS cells, grown in 2xYT media at 37 °C, and expressed at OD 0.8 for 4 h with 0.5 mM IPTG. Cells were harvested and resuspended in buffer-A [50 mM HEPES pH 8.0, 40 mM imidazole pH 8.0, and 10% glycerol along with protease inhibitors (1 mM PMSF, 1 μg/ml pepstatin A, and 1 μg/ml leupeptin)] containing 1 M NaCl (A1000). To purify full-length GyrA and GyrB constructs, cells were thawed and lysed by adding lysozyme (2 mg/mL), incubating on ice for 30 mins, and then passing through a HisTrap-HP 5 mL column. After washing with 10 column volumes (CV) of Buffer-A1000, and further with 10 CV Buffer-A containing 150 mM NaCl (A150), protein was eluted directly onto an A150-equilibrated HiTrap-Q 5 mL column. The column was washed with 10 CV of buffer A150, and protein was eluted using a gradient of A150 and A1000. Peak fractions containing the protein were collected, and the His-SUMO tag was removed via overnight digestion at 4 °C using SUMO protease. The mixture was then dialyzed against Buffer A1000 to remove excess imidazole. After tag removal, the cleaved tag and protease were separated from the protein by re-passing the solution over a HisTrap column in Buffer A1000, and the flowthrough was concentrated. The protein was further purified by loading onto a HiLoad Superdex-200 column equilibrated with a buffer containing 500 mM KCl, 50 mM HEPES pH 8.0, and 10% glycerol (S500). Peak fractions, assessed using SDS-PAGE, were collected, concentrated, and adjusted to 20% glycerol to act as a cryoprotectant prior to freezing in liquid nitrogen.

### Plasmid DNA preparation with different DNA supercoiling status

Negatively supercoiled DNA was prepared by incubating with gyrase (5-fold excess relative to the DNA molarity) in 30 mM Tris-HCl (pH 7.8), 200 mM potassium glutamate, 1 mM TCEP, 10 mM Mg(OAc)_2_, 10% glycerol and ATP 5 mM at 37 °C for 1 hour. The gyrase was inactivated at 65 °C for 20 min. Positively supercoiled DNA was prepared by incubating with 9°N Reverse Gyrase (5 units/μg DNA, M0200, provided by New England Biolabs) in 35 mM Tris-HCl (pH 7.5 at 25 °C), 24 mM KCl, 4 mM MgCl_2_, 2 mM DTT, 1.75 mM spermidine, 0.1 mg/ml BSA and 6.5% glycerol at 80 °C for 30 min. In both reactions, T5 exonuclease (5 units/μg DNA, NEB, M0363) was added to the mixture and incubated at 37 °C for 30 min to digest the remaining nicked DNA. Nicked DNA was prepared by BsmI (NEB, R0134S) (2 units/μg DNA) at 65 °C for 1 hour. All plasmid DNA were purified using PCR spin column (Qiagen, 28104).

### Microscopy and data acquisition for single-molecule assays

Microscopy was conducted using a Nikon Eclipse Ti microscope equipped with a custom prism-type TIRFM module, controlled by home-built software (smCamera 2.0) (28). A Nikon 60X/1.27 NA objective (CFI Plan Apo IR 60XC WI) was utilized. Illumination was provided by solid-state lasers (Coherent, 641 nm), which were combined and coupled to an optical fiber. Emission signals were collected through long-pass filters (T540LPXR UF3, T635LPXR UF3, T760LPXR UF3) and a custom laser-blocking notch filter (ZET488/543/638/750M) from Chroma. Images were captured with an electron-multiplying charge-coupled device (EMCCD; Andor iXon 897).

### CRISPR-Cas9

#### dCas9-gRNA complex preparation

For gRNA assemble, crRNA (IDT) and tracrRNA (IDT, 1072532) were diluted in Nuclease-Free IDTE buffer (supplemented with tracrRNA) and mixed at 1:1 ratio in Nuclease-Free water (Final conc. 10 μM). The mixture was heated at 95 °C for 5 min and cooled to room temperature (1 °C/min). The dCas9-gRNA complex was assembled by mixing guide-RNA and dCas9 protein (IDT, 1081066) at a ratio of 2:1 in Cas9 dilution buffer (30 mM HEPES, 150 mM KCl, pH 7.5). RNA sequences are available in Supplementary Table S2.

#### Single-molecule fluorescence imaging and quantification

Detailed methods of smFRET data acquisition and analysis were described previously(29). For the DNA unwinding assay, 200 pM of Cy3-, Cy5- and biotin-labeled plasmid DNA was immobilized on the polyethylene glycol-passivated flow chamber surface (Nano Surface Sciences) using NeutrAvidin-biotin interaction. All the imaging were carried out at room temperature in Cas9-gRNA reaction buffer supplemented with oxygen scavenging reagents and trolox (20 mM Tris-HCl pH 8, 100 mM KCl, 5 mM MgCl_2_, 5% (v/v) glycerol, 0.2 mg/ml BSA, 1 mg/ml glucose oxidase, 0.03 mg/ml catalase, saturated Trolox (> 5 mM) and 0.8% (w/v) dextrose). All movies were recorded with a 50 ms time resolution. Before the addition of the dCas9-gRNA complex, short movies of the first 10 frames under Cy3 excitation and the last 10 frames under Cy5 excitation were recorded at 10 different imaging views. Following the addition of 10 nM dCas9-gRNA complex with a mismatch on PAM-proximal region, the same format of short movies was captured at each of the following time points: 1, 2, 3, 4, 7, 10, 13, 20, 30, 45 and 60 minutes. All data points were used for fitting, but the plot was trimmed to exclude the data points of 30-, 45- and 60-min. Recording 10 movies required approximately 20 seconds and yielded data from around 5000 molecules. In the case of dCas9-gRNA with three consecutive mismatches on PAM-distal region, the movies composed of initial and last 10 frames under Cy5 excitation and 980 frames under Cy3 excitation were recorded from five different fields of view.

#### DNA unwinding kinetics analysis

The first 10 frames were used to calculate FRET efficiency, and the last 10 frames were used to filter the fluorescent signal from inactive or missing acceptor. Five frames from third to seventh frames of each molecule’s FRET trajectories were selected as data points to construct FRET histograms. The FRET histograms were fitted using a sum of three Gaussian functions. The intersection points of the Gaussian curves were determined to divide the population into two groups by FRET. For dCas9-gRNA with a single mismatch on PAM-proximal region, molecules exhibiting FRET values between 0.14 and 0.47 were classified as DNA unwound by the dCas9-gRNA complex. Those with FRET values exceeding 0.14 were categorized as total DNA molecules. In the case of dCas9-gRNA with three consecutive mismatches in PAM-distal region, those with FRET values exceeding 0.18 were categorized as total DNA molecules and those with FRET between 0.18 and 0.51 were classified as DNA unwound. For the dwell time analysis, the FRET traces with dynamics were collected. The FRET dynamics as a function of time was characterized into two states using Hidden Markov Model and the dwell times for each state were collected by ebFRET (30).

### MutS

#### Preparation of fluorescently labeled MutS

*E. coli* MutS was cloned with a N-terminal hexa-histidine and sortase (LPETG) tag into a pET expression plasmid. MutS was expressed in BL21-AI cells with 1 L of culture. The culture was grown at 37 °C until it reached an OD600 of 0.3. The temperature was lowered to 16 °C and 0.2 % L-(+)-Arabinose was added. After 30 minutes 0.05 mM IPTG was added, and the culture grew for 16 hrs. The cells were collected and resuspend in Buffer A (25 mM HEPES pH 7.8, 10% glycerol, 0.8 μg/mL pepstatin, 1 μg/mL pepstatin and 87 μg/mL PMSF) containing 300 mM NaCl and 20 mM imidazole. The resuspended cell pellet was flash frozen and stored at -80 °C. For the purification of MutS, all steps were carried out at 4 °C. The cell pellet was thawed and lysed by sonication. The lysate was clarified by centrifuging at 120,000 x g for 1 h at 4 °C. The supernatant was then applied to a 3 mL Ni-NTA (Qiagen) column. The Ni-NTA column was washed with 10 column volumes of Buffer A containing 800 mM NaCl and 20 mM imidazole followed by 5 column volumes of Buffer A containing 90 mM NaCl and 20 mM imidazole. The Ni-NTA column was then eluted directly onto a tandem 3 mL Heparin-Sepharose (Cytiva) column with 5 column volumes of Buffer A containing 90 mM NaCl and 200 mM imidazole, washed with 10 column volumes of Buffer B (25 mM HEPES pH 7.8 and 10% glycerol) containing 100 mM NaCl and eluted with a 20 mL linear gradient of Buffer B containing 100 mM to 1 M NaCl. Peak fractions were analyzed by PAGE and combined. The combined MutS fractions were fluorescently labeled using a sortase transpeptidase. A total of 13 nmol of MutS was combined with 52 nmol of purified S. aureus sortase 5 M, and 130 nmol of purified Cy3-CLPETGG labeled peptide. The labeling reaction was carried out at 4 °C for 1 hr in Buffer B containing 10 mM CaCl_2_ and 300 mM NaCl. The reaction was then stopped by the addition of 20 mM EGTA. The excess label was removed by passing the reaction mix through a 10K MWCO Zeba spin desalting column (Thermo Scientific) followed by application onto a 1 mL Heparin-Sepharose column. Bound material was step eluted with Buffer C (25 mM HEPES pH 7.8, 600 mM NaCl, 10% glycerol, 0.1 mM EDTA, 1 mM DTT, 0.8 μg/mL pepstatin, 1 μg/mL pepstatin and 87 μg/mL PMSF), which removed the Sortase protein and concentrated the MutS. Peak fractions were dialyzed into storage buffer (25 mM HEPES pH 7.8, 150 mM NaCl, 20% glycerol, 0.1 mM EDTA and 1 mM DTT), flash frozen and stored at -80 °C. We determined a 48% MutS monomer labeling efficiency.

#### Single-molecule fluorescence imaging and quantification

For Must binding assay, Cy5-labeled plasmid DNA containing one GT mismatched base pair was immobilized on the polyethylene glycol-passivated flow chamber surface using NeutrAvidin-biotin interaction. Cy3 labeled *E. coli* MutS (2 nM) was incubated with 1 mM of ADP or ATP in MutS reaction buffer (20 mM Tris-HCl, pH 7.5, 5 mM MgCl_2_, 100 mM potassium glutamate, 0.1 mM DTT, 0.2 mg/mL Acetylated BSA (Invitrogen, AM2614) and 0.0025% Tween 20 (BioRad, 170-6531)) supplemented with oxygen-scavenging system (2.5 mM 3,4-Dihydroxybenzoic acid (PCA) (Sigma, 37580-25G-F) and 50 nM recombinant protocatechuate 3,4-Dioxygenase from bacteria (rPCD or PCD) (OYC, 46852004)), and saturated Trolox (> 5 mM). All movies were recorded at room temperature. In the presence of ADP, long movies consisting of the first 10 frames under Cy5 excitation, followed by 980 frames under Cy3 excitation, and 10 frames under Cy5 excitation were recorded at 3 different imaging views with a time resolution of 200 ms. Under ATP condition, long movies consisting of the first 10 frames under Cy5 excitation, followed by 1480 frames under Cy3 excitation, and 10 frames under Cy5 excitation were recorded at 3 different imaging views with a time resolution of 50 ms. The first 10 frames were used to select DNA.

#### Enzyme kinetics analysis

The dwell times were collected manually using custom MATLAB scripts. In ATP condition, the dwell times of high FRET (*t*_SB_) both followed by low FRET and not followed were collected manually using custom MATLAB scripts. The dwell time was plotted in 1-CDF plot and fitted to single exponential decay function (equation 1, where *t* is the dwell time, *y*_0_ is baseline offset and *A*_1_ is initial value of 1-CDF at *t*=0) to obtain the rate constant (*k*).

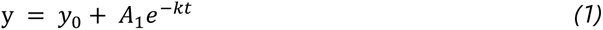

This rate constant is sum of two rate constants, *k*_-1_ (dissociation from mismatch region) and *k*_2_ (conformational change from initial binding to sliding clamp) (equation 2).

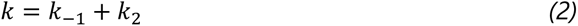

The branch ratio is defined as the number of sliding clamp formations (N_2_) divided by the number of specific binding events not followed by sliding clamp (N_-1_). By setting the equation 3, the *k*_-1_ and *k*_2_ were determined.

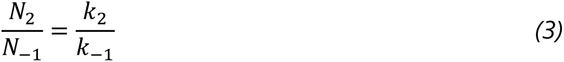

All experiments were performed more than three times, and the error bar represents the standard error.

## RESULTS

### Site-specific labeled plasmid DNA is an optimal sample for smTIRF experiment

Our aim was to achieve site-specific labeling of the circular DNA with fluorophores and biotin so that we can introduce different supercoiling states while taking advantage of FRET analysis of surface-tethered DNA molecules for a detailed exploration of DNA-protein interactions and conformational changes. To site-specifically label the circular DNA, we employed a nicking enzyme-based internal labeling strategy (27). Initially, we cloned pUC19 plasmids to carry two recognition sites for the nicking enzyme BbvCI, approximately 50 bp apart from each other. After nicking the two sites, the small fragment thus produced was evicted by elevating the temperature and replacing it with a single-stranded oligonucleotide labeled with fluorophores and/or biotin by slowly cooling the mixture. Following ligation of the nicks, we successfully generated covalently closed circular DNA with labels (Figure 1A). Through optimization at each step, we retrieved 40% of the initial plasmid DNA, sufficient for generating samples tailored for single-molecule experiments. We quantified the replacement efficiency, assessing how effectively the labeled single-stranded oligonucleotide replaced the native strand. By comparing the fluorescence intensity of the single-stranded oligonucleotide and circular DNA at equal concentration, we determined that approximately 60% of retrieved plasmid DNA had fluorescent labels (Supplementary Figure S1). For further modifications, we repeated this process on the opposite strand. At the last step, we removed all the remaining single stranded oligonucleotide and non-ligated byproduct by treating the mixture with T5 exonuclease (Figure 1B). During the nicking and ligating steps, the supercoiling tension was released, generating relaxed circular DNA. And we subsequently treated the plasmid DNA with gyrase and reverse gyrase to generate negatively and positively supercoiled DNA, respectively (Figure 1C).

**Figure 1:**
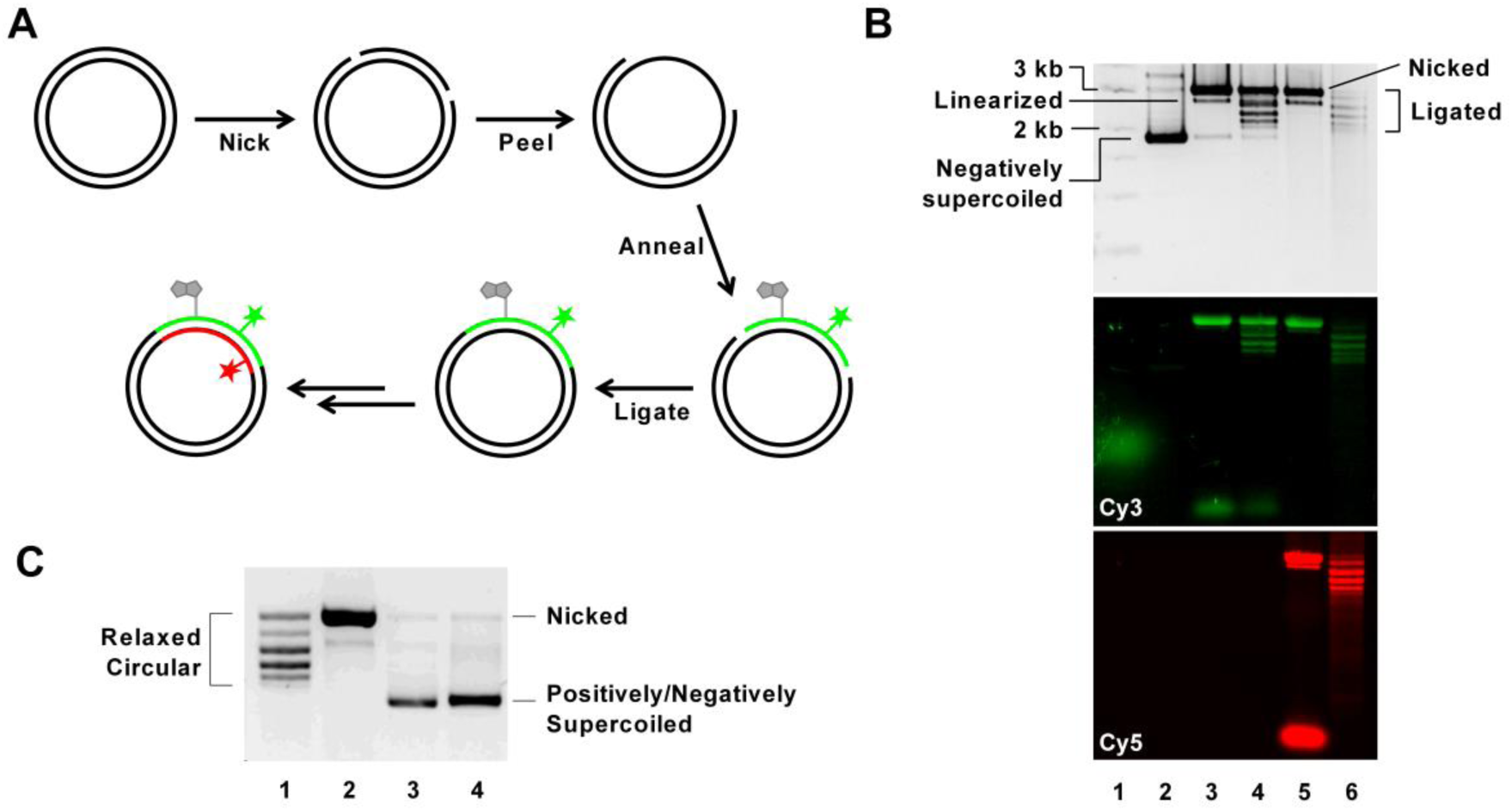
(A) Schematic representation of the plasmid preparation method. A plasmid (pUC19) is modified to carry two BbvCI recognition sites. The plasmid was prepared from E. coli after transformation. Two nicks are introduced into the plasmid using a nicking enzyme recognizing BbvCI sites. The small ssDNA fragment was peeled off at 95 °C and was replaced by a labeled ssDNA of the same size. The nicks are ligated at the last step. The same process can be repeated on the other strand to introduce further modifications. (B) Gel electrophoresis images showing plasmid samples at each preparation step. Lane 1: DNA ladder. Lane 2: Native plasmid isolated from E. coli. The plasmid is negatively supercoiled and migrates faster than a linear DNA of equivalent molecular weight. Lane 3: Plasmid nicked at two sites on one strand and annealed with an oligo with Cy3 and biotin modifications. The nicked plasmid migrates slower than the linear byproduct. The second panel shows Cy3 signal, confirming the insertion of the Cy3-labeled oligo. Lane 4: Ligated plasmid displaying discrete bands after ligation. Lane 5: Plasmid nicked at two sites on the opposite strand of the product in Lane 4 and annealed with a Cy5-labeled oligo. The bottom panel shows Cy5 signal, indicating the successful insertion of the Cy5-labeled oligo. Excess Cy5-labeled oligo appears as the bottom-most band. Lane 6: Plasmid after final ligation, with nicked DNA removed by T5 exonuclease treatment. (C) A gel electrophoresis image showing Cy5-labeled plasmid samples of different supercoiling states. Lane 1: Relaxed circular DNA, Lane 2: Nicked DNA, Lane3: Positively supercoiled DNA, Lane 4: Negatively supercoiled DNA. Gel shows Cy5 signal.

In subsequent experiments, we scrutinized the condition for immobilization of plasmid on surface using TIRF microscopy. By incubating 200 pM of DNA for 5 min at room temperature, we successfully immobilized the DNA molecules with a suitable areal density (∼480 spots per imaging field of view) on polymer-passivated surface using biotin-NeutrAvidin interaction (Supplementary Figure S2). Under these experimental conditions but without NeutrAvidin, the number of nonspecifically adsorbed spots was much smaller (∼55 per field of view frame). These observations demonstrate that our modified plasmid DNA is optimal for smTIRF measurements.

### CRISPR-Cas9: Negative DNA supercoiling enhances R-loop formation in the presence of mismatch

In our initial experiments, we designed DNA constructs to investigate the impact of DNA supercoiling on DNA unwinding by CRISPR-Cas9. We previously developed a smFRET assay to probe DNA unwinding by dCas9-gRNA using short linear DNA molecules labeled with Cy3 (FRET donor) and Cy5 (FRET acceptor) (29). We implemented the same dual labeling system on a circular DNA (Figure 2A). The CRISPR-Cas9 target sequence was introduced to plasmid DNA. Subsequent modifications enabled the site-specific labeling of the target strand and non-target strand with Cy3 and Cy5, respectively. In addition, a biotin was incorporated at a position 22 base pairs away from the protospacer to facilitate the immobilization of the DNA. Before introducing dCas9-gRNA, the protospacer region of the DNA was fully base paired and exhibited high FRET efficiency (Figure 2B). Following the incubation of DNA with dCas9-gRNA, the dCas9-gRNA unwound the protospacer region, displaying lower FRET efficiency. This shift in FRET efficiency reflects the DNA structural changes induced by the interaction of dCas9-gRNA with the protospacer region, providing a dynamic readout of CRISPR-Cas9 unwinding activity on the plasmid DNA construct.

**Figure 2:**
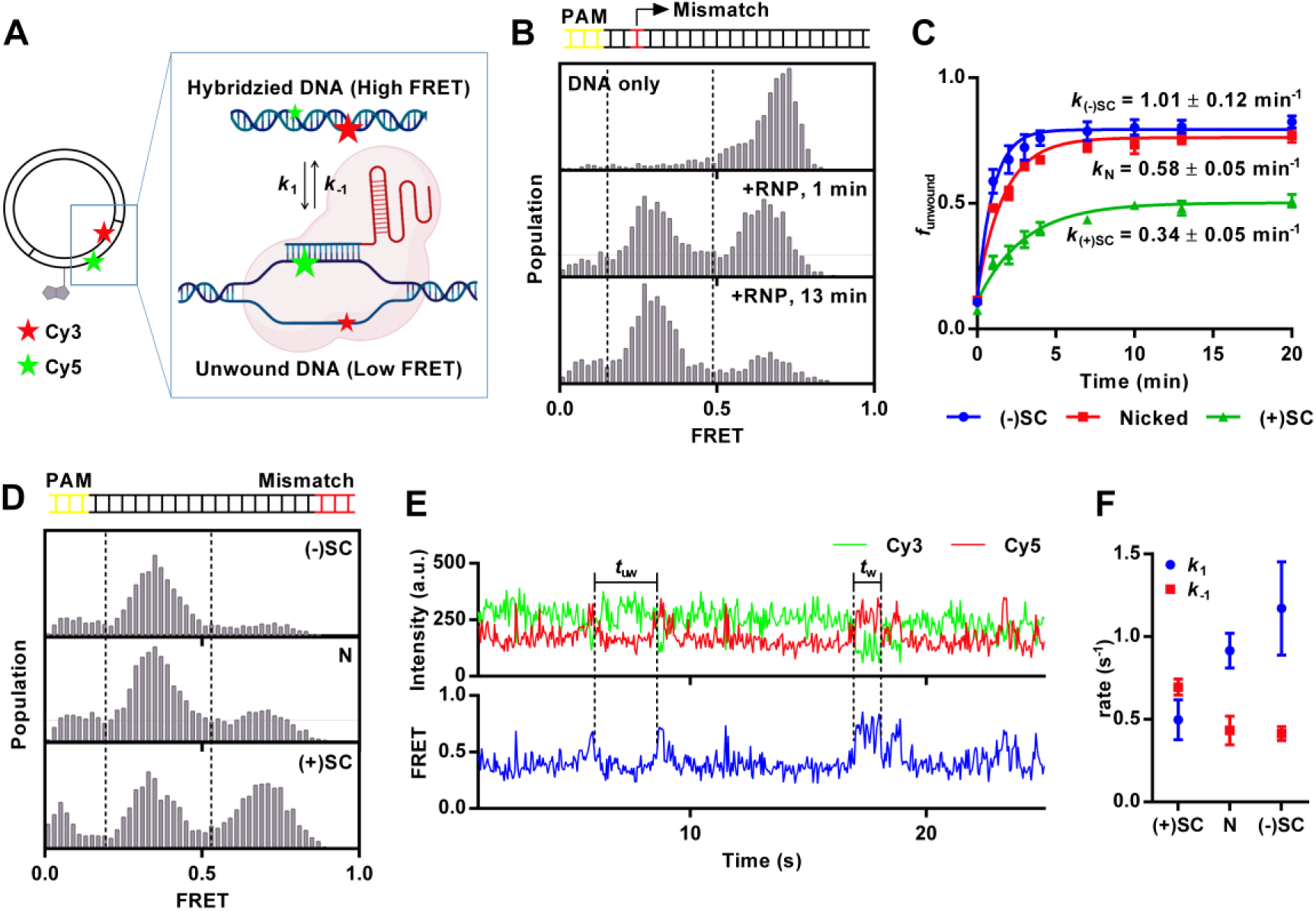
(A) Schematic representation of the experimental design. Plasmid is labeled with a biotin 22 bp away from protospacer, with a donor (Cy3) and an acceptor (Cy5) positioned 10 bp apart on protospacer. The Cy3-Cy5 exhibits high FRET when DNA is fully hybridized, while low FRET indicates protospacer unwinding and R-loop formation by the dCas9-gRNA complex. (B and C) DNA unwinding when the gRNA has a single mismatch 3 bp from the PAM site. (B) FRET histograms of dCas9–gRNA binding to DNA constructs. Nicked plasmid was immobilized on a slide, and 10 nM of dCas9-gRNA was added. High FRET (∼0.6) was observed before the addition of dCas9-gRNA, and after 1 minute of incubation, half of the DNA population shifted to low FRET (∼0.4). After 13 minutes, nearly the entire population exhibited low FRET. (C) Fraction of unwound DNA, funwound, as a function of incubation time for negatively supercoiled ((-)SC), nicked (N) and positively supercoiled ((+)SC) plasmids. Approximately 5000 molecules were recorded over 20 seconds at each time point. Error bars represent standard error for n = 3. (D, E and F) DNA unwinding when the gRNA has three mismatches on 18-20 bp away from the PAM site. (D) FRET histograms of dCas9-gRNA binding to negatively supercoiled ((-)SC), nicked (N) and positively supercoiled ((+)SC) plasmids. (E) Representative single molecule intensity time traces of Cy3 (green), Cy5 (red) and FRET efficiency (blue) showing DNA unwinding dynamics by dCas9-gRNA. (F) Rate constants of the DNA unwinding and rewinding by dCas9-gRNA. The unwinding rate (k1) and rewinding rate (k-1) are indicated as blue circles and red squares, respectively. Over 500 data points are collected from over 100 molecules for each DNA sample. All triplicate data are provided in Supplementary Table S3.

To further investigate the DNA supercoiling effect on DNA unwinding activity of dCas9-gRNA, we induced negative and positive supercoiling in the labeled plasmid DNA using gyrase and reverse gyrase, respectively. We also employed nicked DNA as a “zero supercoiling level” control, achieved through a specific nicking enzyme treatment. The nick site was positioned approximately 1000 base pairs away from the protospacer to ensure that its influence on the CRISPR-Cas9 was limited to the effects of supercoiling relief rather than directly affecting the protein interaction.

To examine the DNA supercoiling effect on off-target binding and unwinding by CRISPR-Cas9, we introduced a single nucleotide mismatch between the target sequence and the gRNA in the PAM-proximal region. Using this dCas9-gRNA complex, the negatively supercoiled DNA displayed the most pronounced shift of FRET populations from high to low, compared to the nicked DNA and positively supercoiled DNA under identical conditions (Figure 2C). To quantify the unwinding activity on three different DNA constructs, we calculated the fraction of unwound state, *f*_unwound_, by dividing the population of low FRET by the sum of low FRET and high FRET populations and plotted *f*_unwound_ over time after addition of 10 nM of dCas9-gRNA. The data were fitted to one-phase association model using the equation 4 to obtain the rate constant (*k*).

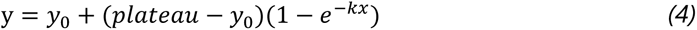

The negatively supercoiled DNA exhibited approximately 2-fold and 3-fold higher rate constant *k* compared to nicked DNA and positively supercoiled DNA, respectively. These results suggest that negative supercoiling enhanced unwinding of target DNA by dCas9-gRNA containing a PAM-proximal mismatch. Additionally, the negatively supercoiled and nicked DNA substrates reached a plateau at 0.79 and 0.76, respectively, indicating higher levels of product accumulation than positively supercoiled DNA, which plateaued at 0.50. Together, these observations suggest positive supercoiling kinetically hinders the unwinding process, likely due to the increased torsional strain on the DNA helix that will impact the efficiency of DNA strand separation (Figure 2C) (31, 32).

Next, we modified the gRNA to introduce three mismatched base pairs between the DNA target sequence and gRNA, spanning positions 18^th^ to 20^th^, counting from the PAM site. Upon addition of the dCas9-gRNA, we measured FRET efficiency in equilibrium (Figure 2D). Negatively supercoiled DNA showed the highest population ratio of DNA unwound (*f*_unwound_ = 0.77), followed by nicked DNA (*f*_unwound_ = 0.67) and then positively supercoiled DNA (*f*_unwound_ = 0.45) (Supplementary Table S3). Additionally, we observed dynamic FRET fluctuations, suggesting continued dynamic DNA conformational changes (Figure 2E). To further analyze the dynamics, we applied hidden Markov models using ebFRET (30), which identified two distinct conformational trajectories characterized by high and low FRET states. We quantified the dwell times for each of these FRET states. The resulting data were transformed into the complement of the cumulative distribution function (1-CDF) and fitted to a first-order exponential decay model, allowing us to calculate the DNA unwinding and rewinding rate constants, *k*_1_ and *k*_-1_. The unwinding rate constant *k*_1_ appeared higher in the order negatively supercoiled DNA> nicked DNA > positively supercoiled DNA. In contrast, the rewinding rate constant *k*_-1_ appeared similar for nicked DNA and negatively supercoiled DNA but was highest in positively supercoiled DNA (Figure 2F). These results indicate that negatively supercoiled DNA exhibits a higher degree of intrinsic unwinding when mismatches were present in the dCas9-gRNA because of faster unwinding and slower rewinding. This demonstrates that negative supercoiling facilitates DNA unwinding by dCas9-gRNA in the presence of mismatches (31, 33, 34).

### MutS: DNA supercoiling stabilizes the interaction between MutS and a mismatched base pair

For an additional demonstration of the versatility of our approach, we extended our study to include *E. coli* MutS, a crucial protein in DNA mismatch repair process (35–37). MutS recognizes and binds to a mismatched base pair while bound to ADP (38). Once bound to a mismatched base pair, MutS exchanges ADP with ATP and undergoes a conformational change, resulting in the formation of a “sliding clamp” (39). Sliding clamp formation is essential for initiating the mismatch repair pathway, as it recruits other repair factors (40–42). Our study focuses on two key aspects of MutS: its binding affinity to the mismatch region, and its transformation into the sliding clamp configuration. We focused on observing MutS behavior with a plasmid DNA containing a GT mismatch. This specific mismatch was chosen due to its known propensity for exhibiting significant MutS binding and sliding clamp formation (43–45). To facilitate this study, the plasmid DNA was labeled with Cy5, positioned 8 base pairs away from the mismatch site (Figure 3A). This labeling position was designed to characterize the interactions with Cy3-labeled MutS via FRET (44, 46). This approach allowed us to observe high FRET efficiency when MutS bound specifically to the mismatched base pair (Figure 3B). Moreover, these MutS binding events were often followed by sliding clamp formation in the presence of ATP, characterized by a transition to low FRET (47). Finally, to investigate the effect of DNA supercoiling on MutS activity, we introduced negative and positive supercoiling into the Cy5-labeled, GT mismatch-containing plasmid DNA using the same approach as in our CRISPR-Cas9 experiments. To avoid potential interference from nicks on MutS (48, 49), we used relaxed circular DNA as the control instead of nicked DNA.

**Figure 3:**
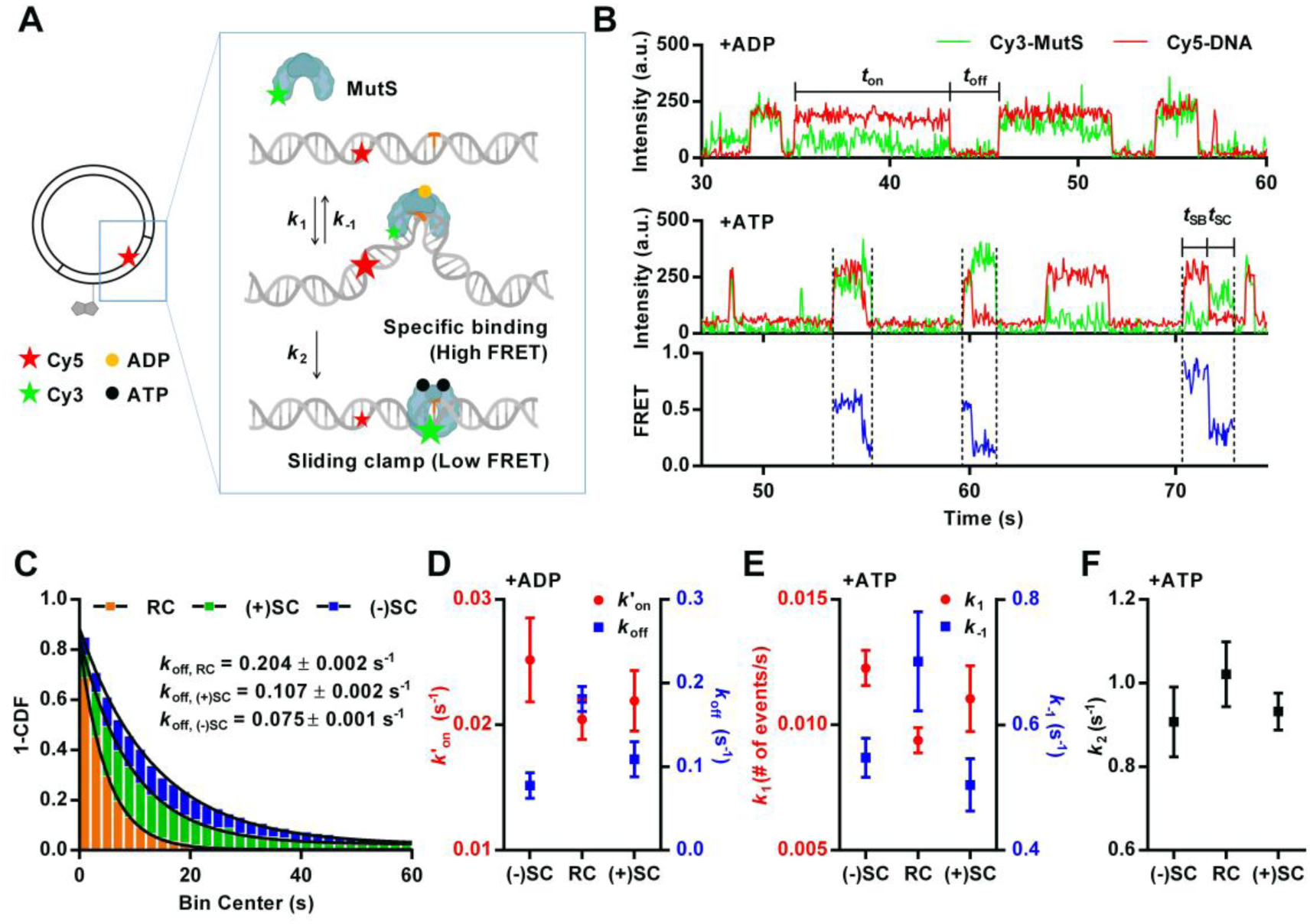
(A) Schematic representation of the experimental design. Plasmid is labeled with a biotin and a Cy5, 26 bp and 8 bp away from a single GT mismatch (colored as orange), respectively. Cy3-labeled, ADP-bound MutS binding to a mismatch causes high FRET between Cy3 on MutS and Cy5 on DNA. MutS undergoes ADP-ATP exchange and forms a sliding clamp, characterized by low FRET. (B) Representative single molecule time traces of Cy3 signal (green), Cy5 signal (red) and FRET efficiency (blue), showing MutS binding to heteroduplex containing a GT mismatch supplemented with 1 mM of ADP (upper panel) and 1 mM of ATP (lower panel). FRET efficiency shows heterogeneities between individual MutS binding events due to the presence of both mono- and di-labeled MutS. (C) MutS dissociation rate (*k*_off_) in the presence of ADP. Binding time (*t*_on_) is collected and transformed into 1-CDF graph. This graph is fitted into single exponential decay function to determine the *k*_off_. Relaxed circular DNA (RC) showed a dissociation rate of 0.204 ± 0.002 s^-1^, positively supercoiled plasmid ((+)SC) showed 0.107 ± 0.002 s^-1^, and negatively supercoiled plasmid ((-)SC) exhibited 0.075 ± 0.001 s^-1^. Approximately 500 events were collected from 200 molecules for each DNA sample. All triplicate data are provided in Supplementary Table S4. (D) Pseudo-first-order association rate (*k*’_on_) and dissociation rate (*k*_off_) of MutS binding to a GT mismatch in the presence of ADP. The error bars represent standard error for n = 3. (E and F) The rate constants, *k*_1_, *k*_-1_, and *k*_2_ in the presence of ATP. The error bars represent standard error for n=4 for negatively supercoiled DNA and relaxed circular DNA, n=3 for positively supercoiled DNA. Each data point collected from approximately 400 molecules.

To determine the binding affinity, we incubated the plasmid DNA with MutS and ADP, which allowed us to exclusively observe high affinity MutS binding without sliding clamp formation (47). As expected, we observed a repetitive pattern of MutS binding to and dissociating from the mismatched region. To quantitatively analyze these interactions, we measured both the dwell time (*t*_on_), representing the duration of MutS binding to the mismatched region as observed through the high FRET value, and the waiting time (*t*_off_), indicating the interval between successive binding events. The collected data was transformed into a 1-CDF and fit to a first-order exponential decay function (Figure 3C). From this analysis, we derived kinetic parameters including pseudo-first-order association rate (*k*’_on_) and dissociation rate (*k*_off_). We observed that negatively supercoiled DNA exhibited a *k*_off_ value approximately 2.7 times smaller than that of relaxed circular DNA. Positively supercoiled DNA also showed slower dissociation, by a factor of 1.4, than that of relaxed circular DNA. The trend for *k’*_on_ between different DNA samples was opposite, though the difference was more subtle. Negatively supercoiled DNA showed 1.2 times faster MutS binding compared to relaxed circular and positively supercoiled DNA has a similar association rate to that of relaxed circular DNA (Figure 3D and Supplementary Table S4). Overall, our data suggest that both positive and negative supercoiling of DNA enhances the stability of the MutS-mismatch interaction in the presence of ADP once binding occurs. However, the initial mismatch binding occurs with similar kinetics on supercoiled DNA and relaxed circular DNA.

Following our observations under ADP conditions, we next examined the impact of ATP on the MutS-mismatch interaction dynamics. Under ATP conditions, we introduced three rate constants to account not only for MutS binding affinity but also its subsequent ATP binding-induced conformational change to form a sliding clamp (Figure 3A). We defined the rate of MutS binding to the mismatch region as *k*_1_, the dissociation rate as *k*_-1_, and the conformational change to the sliding clamp from the initial mismatch-bound state as *k*_2_. Once MutS forms a sliding clamp, subsequent protein dynamics became difficult to analyze because the low FRET state encompasses multiple scenarios, representing both the sliding clamp conformation and the potential movement of MutS away from the mismatch region. The termination of the low FRET state, coupled with the absence of the green signal, signifies either the dissociation of MutS from the DNA or sliding away from the mismatch region beyond the evanescent field of total internal reflection excitation. Therefore, we focused our analysis on sliding clamp formation and not beyond.

To estimate *k*_1_, we counted the number of MutS-mismatch binding events and calculated the number of binding events per second. To determine *k*_2_ and *k*_-1_, we determined two quantities, MutS dwell time on the mismatch (as a specific binding mode, represented by the high FRET state) and the branching ratio between sliding clamp formation and dissociation (*Materials and Methods*). This comprehensive approach allowed us to dissect the dynamic aspects of MutS-mismatch interactions in the presence of ATP. We found that negatively supercoiled DNA exhibited a *k*_1_ value about 1.3 times larger than that of relaxed circular DNA in the presence of ATP (Figure 3E), similar to what we observed in the presence of ADP. We obtained very similar *k*_1_ values between positively supercoiled DNA and relaxed circular. However, negatively and positively supercoiled DNA showed approximately 1.3 times smaller dissociation rate (*k*_-1_) compared to the relaxed circular DNA. The rate of sliding clamp formation, *k*_2_, did not change significantly with either negative or positive supercoiling (Figure 3F). Overall, our data showed that the higher apparent mismatch affinity for supercoiled DNA compared to relaxed DNA is a conserved property even when ATP-induced sliding clamp formation is permitted, and that sliding clamp formation kinetics itself does not depend on the supercoiling state.

## DISCUSSION

Utilizing our plasmid internal DNA modification method, we successfully generated supercoiled plasmid samples with fluorescent, biotin labels and/or a mismatch at specific locations. This advance allowed us to employ smTIRF to investigate the impact of DNA supercoiling on DNA-protein interactions from dozens to hundreds of single molecules in parallel. Our experiments with CRISPR-Cas9 revealed a significant preference for negative supercoiling by the dCas9-gRNA complex during DNA unwinding. This observation aligns with previous studies, such as those using rotor bead tracking (31) and optical tweezer methods (33, 50). For instance, the rotor bead tracking assay demonstrated that negative torque on DNA can induce both half- and fully unwound states of the DNA, required for Cas9 cleavage (51, 52), suggesting a role for supercoiling in facilitating CRISPR-Cas9 activity. Similarly, studies using optical tweezers found that more non-specific binding occurs in negatively supercoiled or strongly stretched DNA (33, 50). However, unlike these previous methods, which investigated DNA binding and unwinding by Cas9-gRNA while altering the DNA supercoiling status, our FRET-based approach measured the real-time kinetics of DNA unwinding at a fixed supercoiling state, providing a more direct analysis of the unwinding process. These findings collectively support the notion that CRISPR-Cas9 exhibits a larger off-target effect on negatively supercoiled DNA, emphasizing the need to consider DNA topology when assessing the specificity and efficiency of CRISPR-Cas9 in gene editing applications.

In our study of MutS, we made a novel observation suggesting that DNA supercoiling could influence MutS binding affinity to the mismatch region. Through dwell time analysis, we discovered that supercoiled DNA stabilized MutS binding to the mismatch region in the presence of ADP. One hypothesis is that DNA supercoiling aids DNA conformational dynamics that includes DNA bending at the mismatch, which may be captured or induced by MutS binding. The mismatched region, having different deformation energy compared to the homoduplex, allows MutS to distinguish the mismatch region by readily bending and kinking the region (38, 53–56). DNA supercoiling further enhances these structural changes, such as kinking or bending (57–61) and the increased ability to deform DNA structure provides a rational for why supercoiled DNA exhibits a longer binding time with MutS (38, 53–56, 62). Since the MutS ATP-induced conformational change into a sliding clamp is coupled with DNA unbending, it was formally possible that DNA supercoiling could affect the efficiency of MutS sliding clamp formation. However, we observed only a subtle difference in the rate of MutS sliding clamp formation between different supercoiling states. These results suggest that DNA supercoiling may play a role in the MutS initial recognition step and/or maintaining the bent state when MutS is bound to DNA but is unlikely to influence the step that result in sliding clamp formation. The conformational changes of MutS includes multiple steps such as DNA unbending and ADP-ATP exchange (39, 63). These multiple steps potentially obscure any effects of DNA supercoiling on DNA unbending by MutS.

Previously, we found that mismatch repair efficiency *in vivo* is highly sequence-dependent, not only influenced by the mismatch itself but also by the adjacent sequence context (45, 62). We also found that MutS binding affinity to various mismatch sequence configurations in vitro correlates with repair efficiency *in vivo* and these differences originate from the altered DNA dynamics of the mismatch-containing sequences (45, 62). Our current findings, showing that MutS binds to the same mismatch differently depending on the supercoiling state, support our previous conclusion that MutS senses the altered properties of the DNA that a mismatched nucleotide induces into the duplex structure, which may explain how it identifies mismatches in different sequence context (45, 62). Further investigations are warranted to unravel the intricate details of how MutS senses these differences.

While our findings provide valuable insights into the effects of DNA supercoiling on CRISPR-Cas9 and MutS interactions, we can enhance our current platform to better study supercoiling dynamics as they occur within living cells. In cells, DNA exhibits local supercoiling that dynamically changes in response to DNA metabolism processes such as transcription and replication (6, 8, 9, 11, 58, 64). In eukaryotes, the DNA is wrapped around histone octamers that produce a slightly negatively supercoiled chromatin structure (65). The relatively unconstrained DNA within chromatin can acquire negative supercoiling up to an upper limit of σ = -0.07 due to transcription *in vivo* (66). The negative supercoiling generated in our plasmid system closely mirrors the supercoiling states typically observed in cells, particularly when considering that *E. coli* gyrase generates a supercoiling density of approximately -0.06 (67). Since the supercoiling effect caused by transcription varies depending on factors such as the distance from the promoter and the strength of the promoter itself (8), future studies could introduce a promoter of interest into the plasmid to better mimic cellular conditions and further explore these effects. Given that transcription-induced supercoiling can diffuse across a distance over 1.5 kb (8), our plasmid (2.8 kb) is well-suited to accommodate and study these effects.

In conclusion, our platform designed to study DNA supercoiling effects in DNA-protein interactions holds promise for investigating various DNA processing enzymes that introduce altered DNA conformations (68) or are affected by DNA flexibility such as transcription factors (69). Additionally, we can use the labeling strategy to produce other substrates, such as mismatch repeat assembly, which may be applied to optical tweezer or other force spectroscopy systems. This versatility expands the potential applications of the platform beyond DNA repair processes, offering a valuable tool for exploring the dynamic interactions of various enzymes with DNA.

## ACKNOWLEDGEMENTS

We would like to thank Dr. Myung Hyun Jo for his training of H. L. on smTIRF. We also would like to thank Nathan Tanner and Jennifer Ong from New England Biolabs for providing purified 9°N Reverse Gyrase. This article is subject to HHMI’s Open Access to Publications policy. HHMI lab heads have previously granted a nonexclusive CC BY 4.0 license to the public and a sublicensable license to HHMI in their research articles. Pursuant to those licenses, the author-accepted manuscript of this article can be made freely available under a CC BY 4.0 license immediately upon publication.

