## Supplementary Data for "A high throughput single molecule platform to study DNA supercoiling effect on protein-DNA interactions"

| 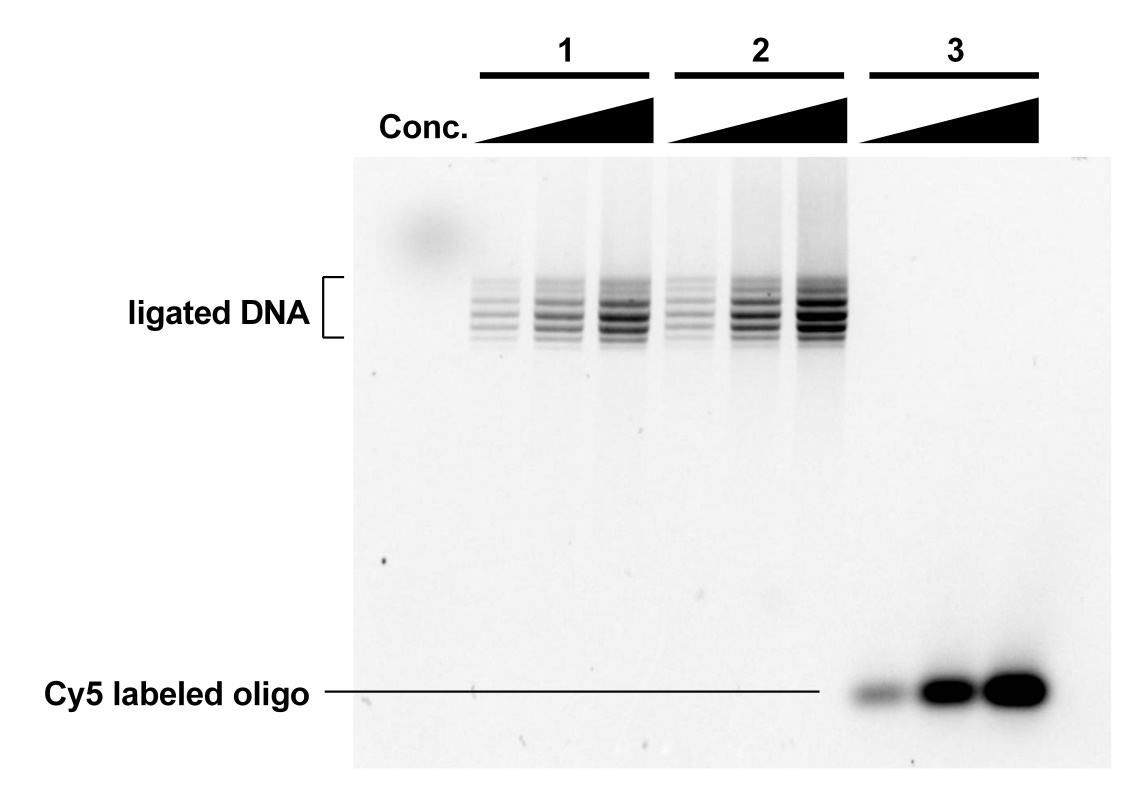 | **Cy5 Intensity** | | | **Cy5 Intensity ratio**  **(ligated DNA : ssOligo)** | | |  |
| --- | --- | --- | --- | --- | --- | --- | --- |
| **Sample** | **Lane 1** | **Lane 2** | **Lane 3** | **Lane 1** | **Lane 2** | **Lane 3** | **Mean** |
| **1** | **6373.75** | **5148.85** | **5046.20** | **0.549** | **0.433** | **0.424** | **0.469** |
| **2** | **9543.82** | **8242.46** | **7469.63** | **0.822** | **0.694** | **0.628** | **0.715** |
| **3** | **11607.08** | **11877.61** | **11903.37** |  |  |  | **0.592** |

**Supplementary Figure S1:** Evaluation of Cy5-labeled Oligo Incorporation into Plasmid DNA. A portion of the plasmid DNA was replaced with Cy5-labeled single-stranded oligo DNA using the strand replacement method described in main text. Two plasmid DNAs were constructed on two separate days. The plasmids and Cy5-labeled oligo were loaded onto a 1% agarose gel at three different concentrations. At each concentration, the Cy5 fluorescence intensity of the ligated plasmid DNA (where the oligo was incorporated) was compared to that of the Cy5-labeled single-stranded oligo to evaluate the efficiency of oligo incorporation into the plasmid. Quantification of the fluorescence intensity was used to calculate the efficiency, and the results are presented in the table below.

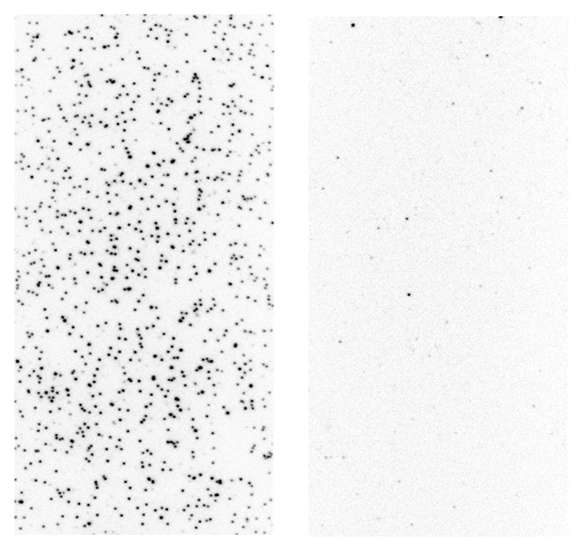
Supplementary Figure S2: Specific binding and non-specific binding of site-specifically labeled plasmids. Cy5 labeled DNA (200 pM) was incubated on the NeutrAvidin treated (left) verse non-treated (right) surfaces for 10 min at room temperature and imaged using Cy5 excitation/emission.

Supplementary Table S1. Single-stranded oligo nucleotides sequence

|  | | DNA sequence |
| --- | --- | --- |
| Rloop | Top | 5’-Phos-TCAGCCCAGCGTCTCATCTTTATACATCAGCAGAGATTTCTGCTGTGCAACC |
|  | Bottom | 5’-Phos-TGAGG**T**TGCACAGCAGAAATCTCTGCTGATGTATAAAGATGAGACGCTGGGC |
| MutS | Top | 5’-Phos-TCAGCT**T**AATACGACTCACTATAGGCCAATACA**G**GAGCTTCATCC |
|  | Bottom | 5’-Phos-TGAGGATGAAGCTCTTGTATTGGCCTATAGTGAGTCGTATTA AGC |

* T: amino-dT labeled with Cy5, T: amino-dT labeled with Cy3, G: GT mismatch, **T**: Biotinylated T

Supplementary Table S2. Crispr RNA sequence

| Mismatch position from PAM | RNA sequence |
| --- | --- |
| 3 | rGrArUrGrUrArUrArArArGrArUrGrArGrArArGrCrGrUrUrUrUrArGrArGrCrUrArUrGrCrU |
| 18-20 | rArGrCrGrUrArUrArArArGrArUrGrArGrArCrGrCrGrUrUrUrUrArGrArGrCrUrArUrGrCrU |

Supplementary Table S3. Rate constants of DNA unwinding and rewinding by dCas9-RNP.

|  | ***k*_1_** | | | | |
| --- | --- | --- | --- | --- | --- |
|  | **Exp 1** | **Exp 2** | **Exp 3** | **mean** | **standard error** |
| **(-)SC** | 1.070 | 0.952 | 1.488 | 1.170 | 0.163 |
| **N** | 0.796 | 0.953 | 0.995 | 0.915 | 0.060 |
| **(+)SC** | 0.629 | 0.391 | 0.471 | 0.497 | 0.070 |
|  | ***k*_-1_** | | | | |
|  | **Exp 1** | **Exp 2** | **Exp 3** | **mean** | **standard error** |
| **(-)SC** | 0.394 | 0.384 | 0.461 | 0.413 | 0.024 |
| **N** | 0.398 | 0.532 | 0.368 | 0.432 | 0.050 |
| **(+)SC** | 0.664 | 0.669 | 0.750 | 0.694 | 0.028 |
|  | ***F*_unwound_** | | | | |
|  | **Exp 1** | **Exp 2** | **Exp 3** | **mean** | **standard error** |
| **(-)SC** | 0.736 | 0.798 | 0.790 | 0.775 | 0.019 |
| **N** | 0.652 | 0.662 | 0.693 | 0.669 | 0.012 |
| **(+)SC** | 0.456 | 0.424 | 0.478 | 0.453 | 0.016 |

Supplementary Table S4. Rate constants of MutS association and dissociation in the presence of ADP.

|  | ***k*_off_** | | | | |
| --- | --- | --- | --- | --- | --- |
|  | **Exp 1** | **Exp 2** | **Exp 3** | **mean** | **standard error** |
| **(-)SC** | 0.0749 | 0.0521 | 0.1049 | 0.0773 | 0.0153 |
| **(+)SC** | 0.1066 | 0.0738 | 0.1462 | 0.1089 | 0.0209 |
| **RC** | 0.2043 | 0.1532 | 0.1852 | 0.1809 | 0.0149 |
|  | ***k*'_on_** | | | | |
|  | **Exp 1** | **Exp 2** | **Exp 3** | **mean** | **standard error** |
| **(-)SC** | 0.0244 | 0.0198 | 0.0313 | 0.0252 | 0.0033 |
| **(+)SC** | 0.0223 | 0.0176 | 0.0259 | 0.0219 | 0.0024 |
| **RC** | 0.0208 | 0.0175 | 0.0230 | 0.0204 | 0.0016 |
|  | ***k*_on_** | | | | |
|  | **Exp 1** | **Exp 2** | **Exp 3** | **mean** | **standard error** |
| **(-)SC** | 1.220E+07 | 9.909E+06 | 1.566E+07 | 1.259E+07 | 1.673E+06 |
| **(+)SC** | 1.115E+07 | 8.776E+06 | 1.295E+07 | 1.096E+07 | 1.209E+06 |
| **RC** | 1.041E+07 | 8.752E+06 | 1.148E+07 | 1.021E+07 | 7.948E+05 |
